# Anti-manic effect of deep brain stimulation of the ventral tegmental area in an animal model of mania induced by methamphetamine

**DOI:** 10.1101/2023.07.02.547148

**Authors:** Roger B. Varela, Suelen L. Boschen, Nathanael Yates, Tristan Houghton, Charles D. Blaha, Kendall H. Lee, Kevin E. Bennet, Abbas Z. Kouzani, Michael Berk, João Quevedo, Samira S. Valvassori, Susannah J. Tye

## Abstract

**Background:** Treatment of refractory bipolar disorder (BD) is extremely challenging. Deep brain stimulation (DBS) holds promise as an effective treatment intervention. However, we still understand very little about the mechanisms of DBS and its application on BD.

**Aim:** The present study aimed to investigate the behavioural and neurochemical effects of ventral tegmental area (VTA) DBS in an animal model of mania induced by methamphetamine (m-amph).

**Methods:** Wistar rats were given 14 days of mamph injections, in the last day animals were submitted to 20 minutes of VTA DBS in two different patterns: intermittent low frequency stimulation (LFS) or continuous high frequency stimulation (HFS). Immediately after DBS, manic-like behaviour and nucleus accumbens (NAc) phasic dopamine (DA) release were evaluated in different groups of animals through open-field test and fast-scan cyclic voltammetry. Levels of NAc dopaminergic markers were evaluated by immunohistochemistry.

**Results:** M-amph induced hyperlocomotion in the animals and both DBS parameters reversed this alteration. Mamph increased DA reuptake time post-sham compared to baseline levels, and both LFS and HFS were able to block this alteration. LFS was also able to reduce phasic DA release when compared to baseline. LFS was able to increase dopamine transporter (DAT) expression in the NAc.

**Conclusion:** These results demonstrate that both VTA LFS and HFS DBS exert anti-manic effects and modulation of DA dynamics in the NAc. More specifically the increase in DA reuptake driven by increased DAT expression may serve as a potential mechanism by which VTA DBS exerts its anti-manic effects.

## 1. Introduction

Bipolar disorder (BD) is a major psychiatric disorder characterized by intermittent and interspersed episodes of mania/hypomania and depression. Although manic episodes are the clinical hallmark for this disorder, BD patients typically spend three times as long with depression as mania/hypomania (1). In fact, bipolar depression is considerably more persistent than non-bipolar depression (2). Lithium, the gold standard mood stabilizer, is used for treatment of bipolar depression (3). However, about two thirds of BD patients may not respond to mood stabilizer treatment and, refractory bipolar treatment is even more challenging due the potential risk of mood shift (4, 5).

The treatment of psychiatric disorders is challenging, as it involves dysfunctions of multiple brain regions and often presents overlapping symptoms and pathologies. Consequently, there is an increasing focus on how the dysfunction of specific brain circuits can lead to specific transdiagnostic psychiatric symptoms (6). One of the more important and studied brain circuits involved in BD is the mesoaccumbens dopaminergic pathway with dopamine (DA)-containing neuronal cells in the ventral tegmental area (VTA) that directly innervate the nucleus accumbens (NAc) (7, 8). Dysregulation of mesoaccumbens dopaminergic transmission in the NAc is often implicated in disorders of motivated behaviour, including BD (9).

For those mood disorders patients who do not respond to any available antidepressant treatments, deep brain stimulation (DBS) holds promise as an effective treatment intervention (10). Recently Drobisz and Damborská (11) compiled evidence reports demonstrating that stimulation of different brain regions can exert antidepressant effects in treatment-resistant depression patients. Although suggesting promising results, DBS for treatment-resistant depression carries many issues that need to be addressed. Very little is understood about the mechanisms of DBS for treatment of depression and moreover, the application of DBS on BD is even less described, as few clinical DBS studies have included BD patients (12). The main concern for the application of DBS on bipolar patients is the observation of treatment associated adverse events in isolated cases (e.g., changes in mood, induced hypomania state). However, the studies about its applicability are still controversial as noted in the current literature, highlighting the need for further studies on the effects of different patterns of stimulation in this condition (13).

It is known that the electrical stimulation parameters have an important role in the mechanism of action of DBS. High frequency stimulation (HFS) (130–185 Hz) induces neuronal depolarization and generates action potentials by opening local voltage-gated sodium channels (14, 15). However, HFS may have a more pronounced inhibitory effect than low frequency stimulation (LFS) under 100 Hz (16). This is thought to occur due to a ‘depolarization block’, where repeated stimulations inhibit sodium ion driven action potentials (17). Also, HFS could have different neural activity effects depending on targeted excitatory or inhibitory neurons (18). Alternatively, Friedman and colleagues (19) designed a programmed LFS based on the difference of VTA neuronal firing patterns in a genetic rat model of depression and control animals. This programmed LFS consists of bursts containing numerous spikes, mimicking and resetting the natural VTA firing configuration, thereby potentially alleviating depressive symptoms in rats (16, 20).

Animal models are important tools to investigate DBS therapeutic mechanisms and applications as they provide a means to control variables that are not easily possible in human studies. Under these conditions, new targets and patterns of stimulations can be tested to refine the functionality of this interventive technique (11, 21). In fact, a genetic animal model suggests that increased firing rate of dopaminergic neurons in the VTA is a key neuronal mechanism for the manic-like behaviors (22). Methamphetamine (m-amph) administration in rats is an animal model of human mania with face, construct and predictive validities, since it can induce manic-like behaviours (including hyperactivity) and responds to mood stabilizers treatment (23). M-amph increases extracellular levels of DA through multiple mechanisms, mindful that dopamine is a critical element in bipolar pathophysiology (24) and the NAc is particularly affected due to its high concentration of DA terminals (25, 26).

There are several potential mechanisms through which DBS of the mesoaccumbens dopaminergic pathway could correct aberrant DA signaling in the NAc that is characteristic of mania. In line with the depolarization block hypothesis of DBS, inhibiting DA release from the VTA to the NAc should reduce the synaptic concentrations of DA in the NAc induced by m-amph. Alternatively, DBS applied to this region could also reset the natural firing configuration, modulating aberrant activity in the mesolimbic DA circuit, normalizing synaptic DA concentrations in the NAc and consequent abnormal behavior. Therefore, the present study aimed to investigate the effects of VTA DBS (both LFS and HFS) in an animal model of mania induced by m-amph to better describe the effects of DBS on mania states, and potential application for BD treatment.

## 2 Material and methods

### 2.1 Animals

The subjects were adult male Wistar rats (N=15-20/group, weighting 250–350 g) obtained from both the Mayo Clinic (US) or Animal Resources Centre (AUS) breeding colonies. The animals were housed two or five per cage (for behaviour and DA quantification, respectively), with food and water available *ad libitum* and maintained on a 12-h light/dark cycle (lights on at 6:00 a.m.) at a temperature of 22±1 °C. All experimental procedures were performed in accordance with, and with the approval of either the Mayo Clinic or The University of Queensland Institutional Animal Care and Use Committees, respectively. All experiments were performed at the same time during the day to mitigate circadian variations.

### 2.2 Experiment 1: Effects of VTA DBS on mania-like behaviour

After a week of acclimatization, rats received 14 days of either saline (NaCl 0.9%), or m-amph (1mg/kg) treatment via intraperitoneal (I.P.) injections (Sigma-Aldrich, USA). On day 10, animals were anesthetized with isoflurane (1–3%) for stereotactic surgery. A 1.5–2 cm incision was made, and a small hole was broached in the skull at the site corresponding to the target. Twisted bipolar stainless steel electrodes (10 mm long, 0.075 mm in diameter; Plastics One, Virginia, USA) were unilaterally implanted into the VTA (from bregma: antero-posterior (AP) -6.8mm, medial-lateral (ML) +0.6mm and, dorsalventral (DV) −8.6 mm) (27). Stimulating electrodes and four screws were mounted to the skull with dental cement. A control group was not subjected to stereotactic surgery and had no electrode implanted. Unilateral implantation was applied since a previous study reported that unilateral stimulation of this brain region was sufficient for clinical outcomes (28).

At the 15^th^ day, animals received a single I.P. injection of m-amph or saline (challenge), 1 hour and 40 minutes after the injection the animals were submitted to 20 minutes of DBS in two different patterns: Intermittent LFS (Frequency: 10Hz; 2ms pulse width; 2 burst/sec; 300μA; monophasic) or continuous HFS (130Hz; 2ms pulse width; 200μA; monophasic) controlled through Iso-Flex/Master-8 system (AMPI^©^, Israel) (Figure 1A) (n=12-16). Stimulation parameters were based on Gazit et al. (16), mimicking physiological pattern of VTA firing and Kouzani et al. (29), current clinical protocol. Sham animals were submitted to the same surgical procedure, had the electrode implanted and were attached to the stimulation device for the same period as the active DBS groups, but stimulation was turned off. Immediately after the DBS session, animals’ locomotor activity and risk-taking behaviour were evaluated through the open-field test (OFT). For this test, animals were placed in an arena (50x50cm) surrounded by 50 cm high walls. A video camera on the top of the arena was connected to a computer for recording behavioural parameters. The animals were carefully put into the central area and then left to explore the arena for 20 min. Total distance travelled (locomotion) and the time spent in the central area of the arena (risk-taking behavior) were recorded through the EthoVision XT behavioral software (Noldus^©^, Netherlands). Experiment 1 timeline is depicted in Figure 1B. Timepoints for m-amph injection, DBS stimulation and OFT were designed for timing the pharmacokinetics of m-amph with duration of DBS session according to original animal model (23), and Gazit et al. study (16).

**Figure 1:**
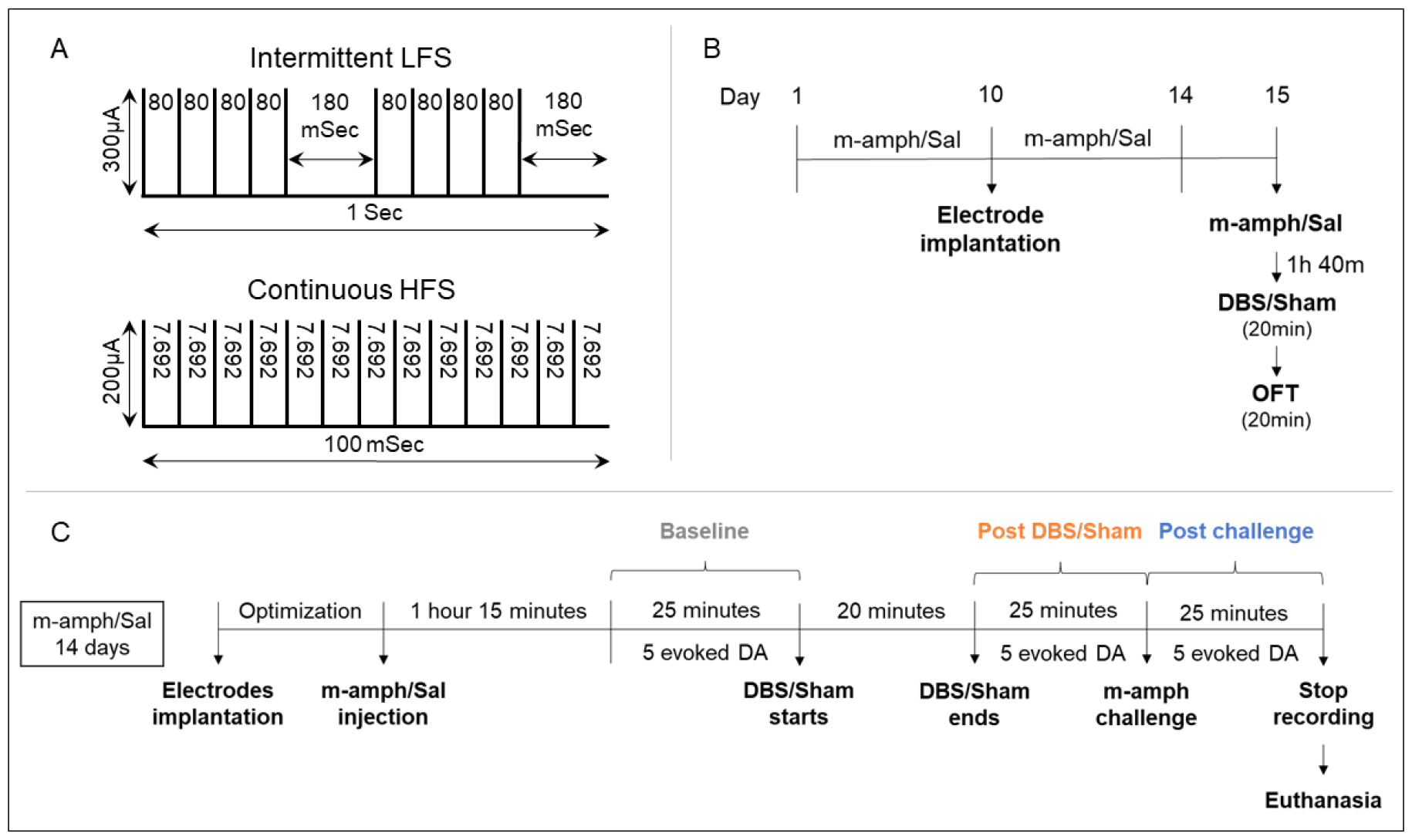
(A) Intermittent low frequency stimulation (LFS): 10Hz stimulation divided in 2 bursts per second, each burst composed of 5 spikes with an inter-spike interval of 80ms and a pause of 180 ms after each burst, according to Gazit et al., 2015. High frequency stimulation (HFS): Continuous stimulation at 130Hz; (B) Experiment 1: Animals received daily intraperitoneal injections of methamphetamine (m-amph) or Saline (Sal) for 14 days. On the tenth day animals were submitted to stereotactic surgery for unilateral implantation of a deep brain stimulation (DBS) electrode in the ventral tegmental area. On the fifteenth day, animals received a single injection of m-amph or Sal and were submitted to 20 minutes of LFS or HFS DBS prior to open field test (OFT); (C) Experiment 2: Animals were submitted to 14 daily injections of m-amph or Sal. On the fifteenth day animals were submitted to fast-scan cyclic voltammetry surgery. After implantation of electrodes and optimization, animals received a single injection of mamph or Sal and had five evoked dopamine (DA) responses recorded before 20 minutes DBS session. Five evoked DA responses were recorded post-DBS session and after m-amph challenge. After FSCV recordings animals were euthanised and brains were removed.

### 2.3 Experiment 2: Effects of DBS on VTA-evoked phasic DA release in the NAc

This experiment was designed to investigate the biological mechanisms underlying DBS effects in significant conditions observed in experiment 1. After a week of acclimatization, rats received 14 days of either saline or m-amph treatment. On the 15^th^ day, animals were submitted to fast-scan cyclic voltammetry (FSCV) analysis for evaluation of VTA-evoked phasic DA release in the NAc. Prior to FSCV surgery, animals were anaesthetised with a 1.5 mg/kg intraperitoneal urethane injection. An incision was made in the midline of the skull to expose bregma. Using bregma as reference, a 1 mm diameter hole was drilled into the skull at the site for the DA recording electrode. The target for the recording electrode was the NAc (AP +1.2, ML +1.5, DV –6.5-8.0 from bregma). The VTA stimulation electrode was implanted according to coordinates described above. A hole was also made for the reference electrode in the opposite hemisphere to the stimulating and recording electrodes. A wireless instantaneous neurotransmitter concentration sensing (WINCS) Harmoni system (Mayo Clinic, USA) was used to generate a voltage-current plot. The location of the recording and stimulating electrodes were optimised by applying stimulation of the VTA (DA-evoked stimulation: 120 pulses; 60Hz; 300-500µA) and recording via the NAc-implanted carbon fiber recording electrode when located at progressively ventral DV coordinates. The recording electrode was lowered to improve the oxidation/reduction signal strength. Adjustments were made until there was a strong signal. Once recording and stimulation sites were optimized, animals received an injection of saline or m-amph (according to pre-treatment).

75 minutes after m-amph/saline injection, five baseline VTA-evoked DA responses were recorded every 5 minutes, period named pre-DBS. Animals were then submitted to VTA DBS (HFS or LFS) or Sham (no stimulation) for 20 minutes (n=4-6). During this time, evoked DA responses were not recorded. After Sham or DBS stimulation, VTA-evoked DA responses were recorded every 5 minutes for 25 minutes, period named post-DBS. After post-DBS recordings, all the animals were submitted to an mamph challenge (1mg/kg) and 5 VTA-evoked DA responses were recorded for confirmation of DA dynamics, period named post-challenge (Figure 1C). The DA peak was determined by averaging the one second time (ten scans) with highest concentration after stimulation. DA reuptake was calculated through T1/2 method which consists in the time for DA concentration to drop to half after peak. The DA recording protocol is demonstrated in the scheme below.

### 2.4 Euthanasia and brain dissection

All the animals were euthanized with I.P. overdose of pentobarbital (150 mg/kg body weight) after the behavior or FSCV protocols. Experiment 2 animals had their brains removed and frozen by placing them on dry ice, followed by storage at − 80 °C for posterior histological and immunohistochemical analysis.

### 2.5 Histological analysis

Frozen tissue sections (10μm) were cut with cryostat (Cryostar NX70, ThermoScientific^©^, USA), mounted on ÜberFrost Slides (InstrumeC^©^, Australia) and immediately fixed using 4% PFA for 10 min. VTA sections were submitted to Nissl staining through cresyl violet for location of the stimulating electrodes. For immunostaining of NAc sections, the tissue section was encircled by using the wax PAP-pen (Daido Sangyo^©^, Japan) and permeabilized with 0.1% Revealit antigen recovery solution (0.1M sodium citrate, citric acid added until pH reaches 6, 0.1M SDS) for 30 min at 40°C using a shaker incubator. Subsequently, sections were washed in PBS for 5 min, and blocking solution for 30 min (0.5 % BSA in 0.05 % saponin, 0.05% sodium azide in 0.1M PBS). Samples were then incubated with the primary antibodies for 4 nights at room temperature protected from light in a humidity chamber. On the fourth day, the sections were washed three times with PBS for 15 min each followed with blocking solution for 5 min. Thereafter, samples were incubated overnight with the secondary antibodies at room temperature. The next day, the sections were washed three times with PBS for 15 min and then coverslipped by using fluorescence mounting medium (Dako^©^, Denmark). Both primary and secondary antibodies were diluted in blocking solution. The primary antibodies used for immunohistochemical analysis included: Mouse monoclonal antibody against Tyrosine Hydroxylase (ImmunoStar Cat# 22941, RRID:AB_572268), rabbit monoclonal antibody against NeuN (Cell Signaling Technology Cat# 24307, RRID:AB_2651140), rat monoclonal antibody against Dopamine Transporter (Millipore Cat# MAB369, RRID:AB_2190413). Goat anti-Mouse IgG Alexa Fluor 488 (Thermo Fisher Scientific Cat# A-11001, RRID:AB_2534069), Goat anti-Rat IgG Alexa Fluor 647 (Molecular Probes Cat# A-21247, RRID:AB_141778) and Donkey anti-Rabbit IgG Alexa Fluor 555 (Molecular Probes Cat# A-31572, RRID:AB_162543) were used as secondary antibodies. Protein expression in the NAc was analysed by semi-quantitative intensity analysis using ImageJ software.

### 2.6 Microscopy and image analysis

All tissue imaging was captured using Zeiss Axio Imager 2 Microscope (https://www.zeiss.com/microscopy/us/products/light-microscopes/axio-imager-2-for-biology.html, RRID: SCR_018876) using a single field of view from the structure of interest, captured using a 20x objective lens, using LED illumination consistent exposure and light settings for all sections belong to a group. The images were then analysed using semi-quantitative immunofluorescent intensity analysis using Fiji software (http://fiji.sc, RRID:SCR_002285) (30) threshold analysis. Identical threshold settings were used between images to ensure consistency. Because DAT is strongly positive in blood vessels, we used a custom written macro to remove the blood vessels from our analysis in these stains (Supplementary Figure 1).

### 2.7 Statistical analysis

Differences among the experimental groups were determined by two-way analysis of variance (ANOVA) followed by Tukey’s post hoc test for OFT. Intragroup differences were determined by one-way ANOVA followed by Tukey’s post hoc test for FSCV recordings and immunohistochemical analysis. In all comparisons, statistical significance was set at p<0.05. Statistical analysis and graphs were performed using Graphpad Prism software, V9.0 (http://www.graphpad.com/, RRID:SCR_002798).

## 3. Results

### 3.1 Open-field test

Behavioral results are demonstrated in Figure 2. Two-way ANOVA revealed a main effect in for m-amph treatment [F(1.95)=5.692, p=0.019] and DBS [F(3.95)=8.421, p<0.0001] in locomotion. Post-hoc analysis revealed that M-amph chronic administration increased locomotor activity when compared to the Saline+control group [Saline+Ctrl vs m-amph+Ctrl: Mean diff= -21878, p=0.0366]. Although electrode implantation in the VTA seems to reduce this m-amph-induced hyperlocomotion [m-amph+Ctrl vs mamph+Sham: Mean diff= 19729, p=0.0847] only LFS and HFS led to a significant reduction [m-amph+Ctrl vs m-amph+LFS: Mean diff= 27388, p=0.0010; m-amph+Ctrl vs m-amph+HFS: Mean diff= 34134, p<0.0001] (Figure 2A). Time spent in the central area of the arena was also assessed as a risk-taking behavior parameter. Although a main effect was observed for m-amph treatment [F(1.95)=7.554, p=0.007], post-hoc analysis did not revealed any relevant difference (Figure 2B).

**Figure 2:**
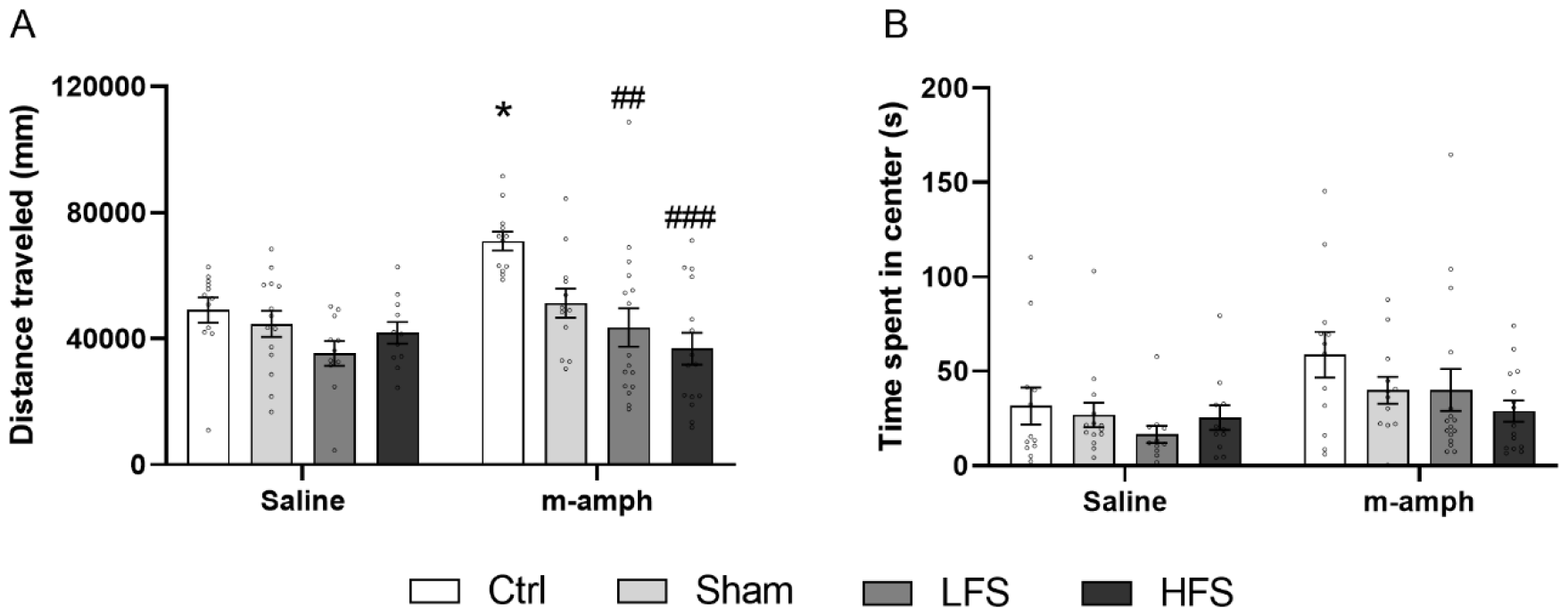
Effects of low frequency stimulation (LFS) and high frequency stimulation (HFS) on travelled distance (A) and time spent in center (B) in the open-field test of rats submitted to the animal model of mania induced by methamphetamine (m-amph). Bars represent mean and standard error according to two-way ANOVA. *=p<0.05 compared to de Saline+Ctrl group; ##=p<0.01 compared to the m-amph+Ctrl group; ###=p<0.001 compared to the m-amph+Ctrl group according to two-way ANOVA followed by Tukey’s post-hoc test.

#### 3.2 VTA-evoked phasic DA release

As depicted in Figure 3, stimulating electrodes were implanted in the VTA region for all animals submitted to the FSCV, except for one (marked in red). Since evoked DA responses were detected in all animals, the misplaced electrode location could have been affected by the brain section used for histological analysis.

**Figure 3:**
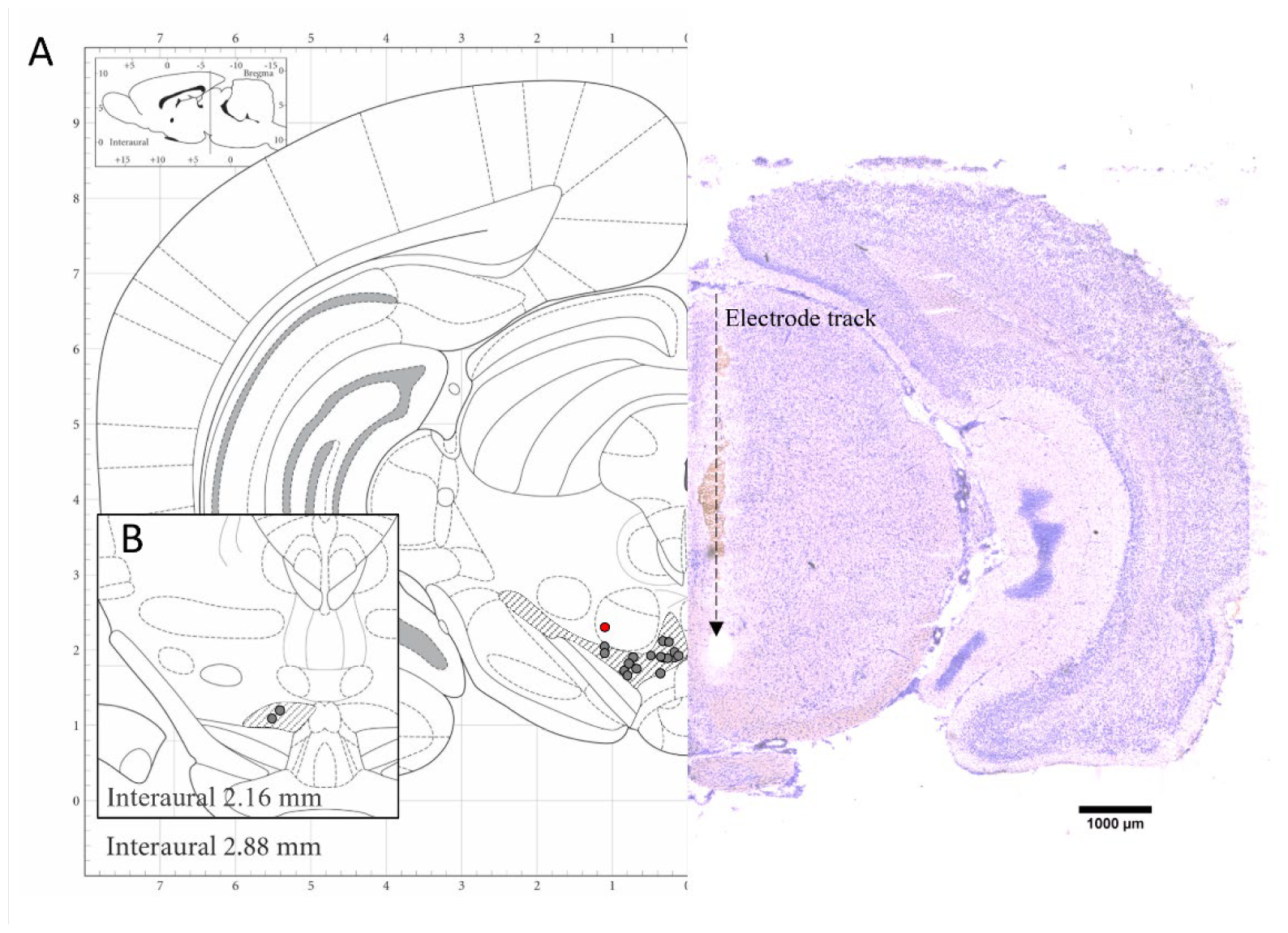
Left: Schematic illustration of coronal rat brain slices at the level of the VTA region (A: interaural 2.16mm; B: interaural 2.88mm). Dots represent implantation sites of stimulating electrodes; red dot represents electrode implantation site outside VTA borders. Image modified from Paxinos & Watson “The Rat Brain in Stereotactic Coordinates (2007). Right: Whole brain mirrored image of VTA DBS animal that was sliced to the site of electrode implantation. Electrode tract mark is visible in this slice.

Results from FSCV recordings are shown in Figure 4. A main effect was observed for Saline+Sham animals regarding DA peak [F(2.85)=28.31, p<0.0001] and reuptake [F(2.84)= 4.516, p=0.0137]. Post-hoc comparison revealed no difference in peak DA concentrations [Mean diff=12.33, p=0.7275] or reuptake [Mean diff=-10.16, p=0.9285] on post-sham compared to baseline, however, mamph challenge increased the DA peak concentration [Mean diff=-97.81, p<0.0001] and reuptake time [Mean diff=-74.99, p=0.0180] in these animals (Figure 4A and 4B). Similar pattern was observed in mamph treated animals, whilst a main effect was observed for DA peak [F(2.72)= 17.05, p<0.0001] and reuptake [F(2.63)=8.037, p=0.0008], post-hoc revealed that sham period did not alter DA peak concentration when compared to baseline [Mean diff=4.267, p=0.8861], but it was increased by m-amph challenge [Mean diff=-43.73, p<0.0001]. In addition, DA reuptake time was enhanced post-sham although not significant [Mean diff=-167.8, p=0.5052], but significantly increased post-challenge [Mean diff=-583.2, p=0.0007] (Figure 4F and 4G). One-way ANOVA also demonstrated a main effect of HFS [F(2.72)= 22.56, p<0.0001] and LFS [F(2.72)= 15.07, p=<0.0001] on DA peak concentration. Multiple comparison analysis shown that, when compared to baseline, DA peak concentration was decreased by both, HFS [Mean diff=18.84, p=0.0202] and LFS [Mean diff=33.80, p=0.0001], but only LFS was able block the effects of m-amph challenge on DA peak concentration [m-amph+HFS baseline vs post-challenge: Mean diff=-26.90, p=0.0006]] (Figure 4K and 4P). Regarding DA reuptake time a main effect of treatment was observed for both m-amph+HFS group [F(2.57)= 16.60, p< 0.0001] and m-amph+LFS group [F(2.72)= 4.878, p= 0.0103]. Post-hoc comparisons revealed that both HFS and LFS did not alter reuptake time when compared to baseline [m-amph+HFS baseline vs post-DBS: Mean diff=-8.806, p=0.7615; mamph+LFS baseline vs post-DBS: Mean diff=-7.558, p=0.8989]. Nor was either treatment effective in blocking the effects of m-amph challenge on reuptake time, as this was significantly increased relative to the observe time to reuptake at baseline [m-amph+HFS baseline vs post-challenge: Mean diff=-66.28, p<0.0001; m-amph+LFS baseline vs post-challende: Mean diff=-49.76, p=0.0136] (Figure 4L and 4Q).

**Figure 4:**
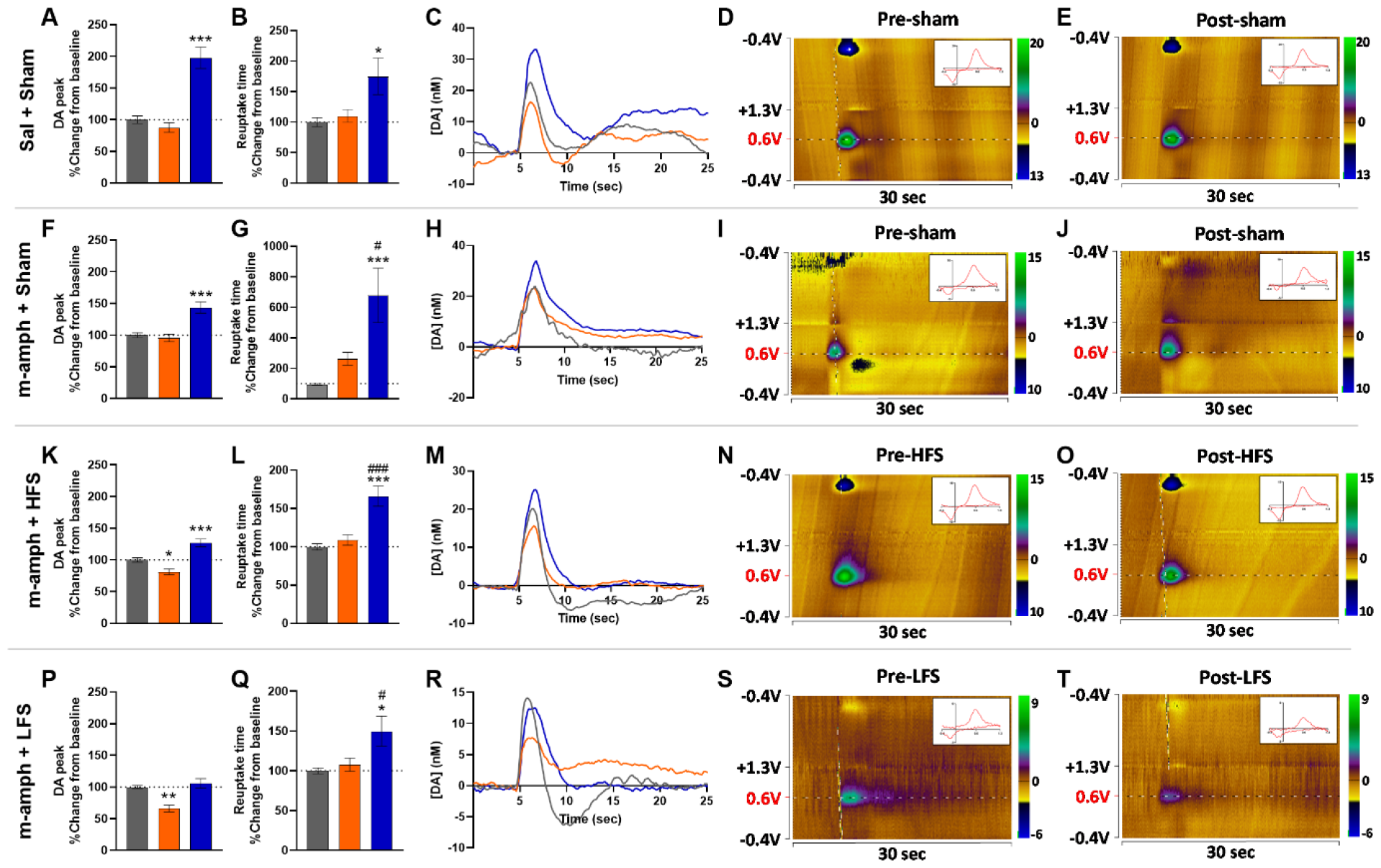
Effects of low frequency stimulation (LFS) and high frequency stimulation (HFS) on nucleus accumbens phasic dopamine (DA) release of rats submitted to the animal model of mania induced by methamphetamine (m-amph). Changes in DA peak concentration (A, F, K and P) and DA reuptake time (B, G, L and Q) are demonstrated in percentage change compared to baseline. Bars represent the mean and standard error. *=p<0.05, **=p<0.01 and ***=p<0.001 compared to the baseline; #=p<0.05, ###=p<0.001 compared to post-Sham/DBS according to two-way ANOVA followed by Tukey’s post-hoc test. Change in evoked DA dynamics are demonstrated in DA concentration over time (C, H, M and R), and pre- and post-DBS/Sham false color plots (D, E, I, J, N, O, S and T) of representative animals.

When comparing DBS effects across groups a main effect on treatment was observed post DBS/Sham period [F(3.99)=4.163, p=0.0080], and post-hoc analysis revealed that only LFS is able to reduce DA peak concentration compared to m-amph+Sham group [Mean diff=29.54, p=0.0054] (Figure 5A). A main effect was also observed for DA reuptake [F(3.99)=13.32, p<0.0001], but post-hoc analysis revealed that both HFS and LFS were able to revert the increase in reuptake time induced by m-amph [m-amph+Sham vs m-amph+HFS: Mean diff=154.6, p<0.0001; m-amph+Sham vs m-amph+LFS: Mean diff=155.8, p<0.0001] (Figure 5C). A main effect on treatment was also observed after challenge for DA peak concentration [F(3.101)=12.53, p<0.0001], post-hoc analysis revealed that m-amph pre-treated animals shown attenuated DA peak concentration comapred to Saline pre-treated animals [Mean diff=54.08, p=0.0054]. Although LFS seemed to decrease DA peak compared to m-amph+Sham group, neigther treatments demonstrated statistically significant reduction [m-amph+Sham vs m-amph+LFS: Mean diff=37.92, p=0.1099; m-amph+Sham vs m-amph+HFS: Mean diff=16.84, p=0.7429] (Figure 5B). Reuptake time also presented a main treatment effect [F(3.101=9.451, p<0.001], multiple comparisons revealed that reuptake time was still increased in m-amph-sham group after challenge [Mean diff=-503.8, p=0.0001], but both HFS and LFS blocked this effect [m-amph+Sham vs m-amph+HFS: Mean diff=512.5, p=0.0002; m-amph+Sham vs m-amph+LFS: Mean diff=529.0, p=0.0001] (Figure 5D).

**Figure 5:**
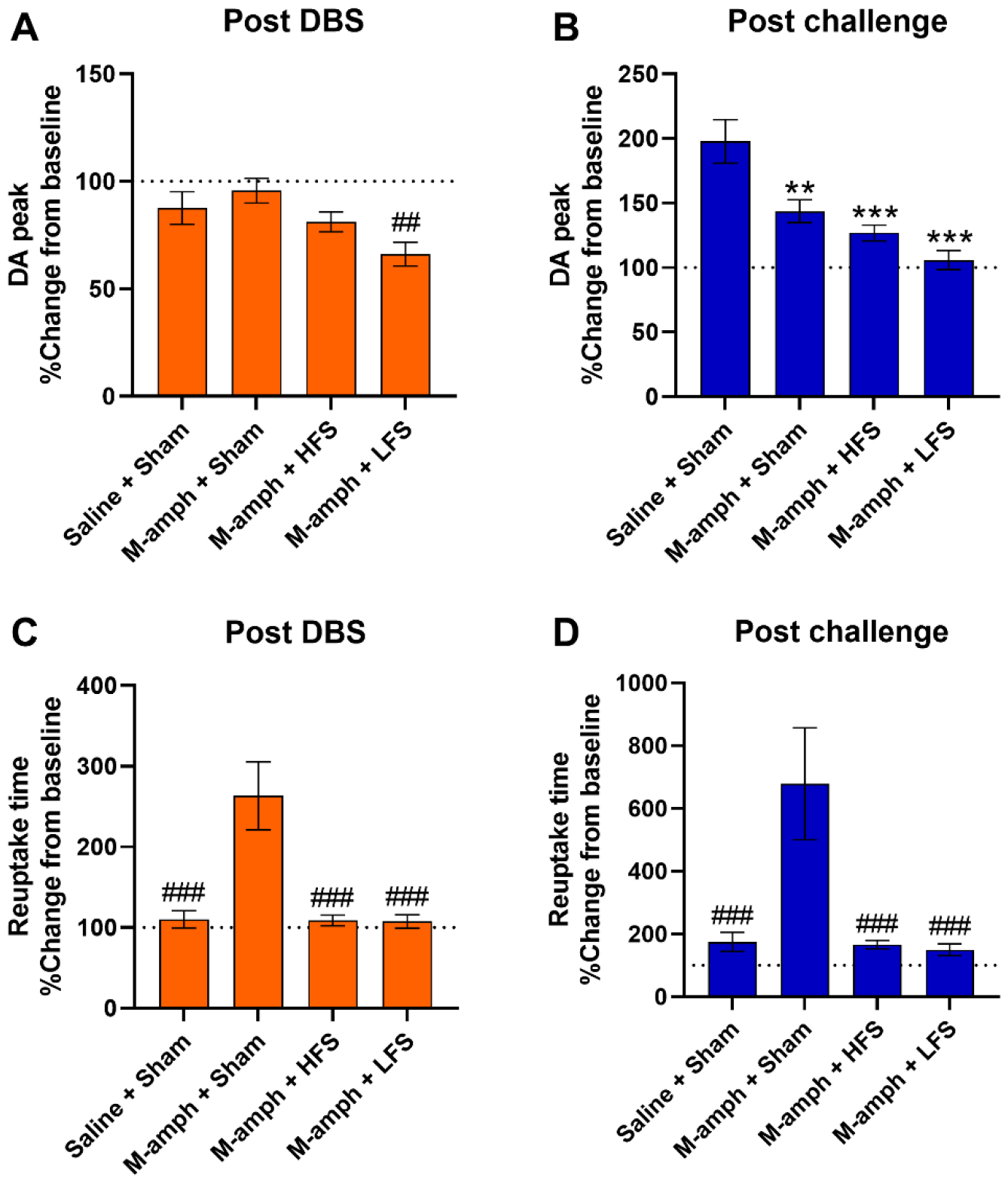
Effects of low frequency stimulation (LFS) and high frequency stimulation (HFS) on dopamine (DA) dynamics across experimental groups. Changes in DA peak (A and B) and reuptake time (C and D) during post-DBS/Sham and post-challenge timepoints (orange and blue bars, respectively). Data is presented in percentage change from baseline, bars represent mean and standard error. **=p<0.01 and ***=p<0.001 when compared to Saline+Sham group; ##=p<0.01, ###=p<0.001 when compared to Mamph+Sham group according to one-way ANOVA followed by Tukey’s post-hoc test.

### 3.3 Immunohistochemistry

Figure 6A demonstrates the expression of DAT, NeuN and tyrosine hydroxylase (TH) in the NAc core and shell, represented as arbitrary intensity units. One-way ANOVA revealed no main treatment effect for NeuN in NAc core [F(3.16)=2.312, p=0.1150] and NAc shell [F(3.16)=0.4935, p=0.6918], neither TH in NAc core [F(3.16)=0.5999, p=0.6243] and NAc shell [F(3.16)=0.4935, p=0.6918]. A main treatment effect was observed in DAT on NAc Core [F(3.16)=4.845, p=0.0138] and NAc Shell [F(3.16)=4.717, p=0.0152]. Post-hoc analysis demonstrates that, when compared to Sham groups, only LFS increased DAT expression in both, NAc core [Saline+Sham vs m-amph+LFS: Mean diff=-1237, p=0.0176; Nac core m-amph+Sham vs m-amph+LFS: Mean diff=-1254, p=0.0313] and shell [Saline+Sham vs m-amph+LFS: Mean diff=-1241, p=0.0155; Nac Shell m-amph+Sham vs m-amph+LFS: Mean diff=-1147, p=0.0482]. No difference was observed in any of the other groups. Representative images from IHC are demonstrated in Figure 6B.

**Figure 6:**
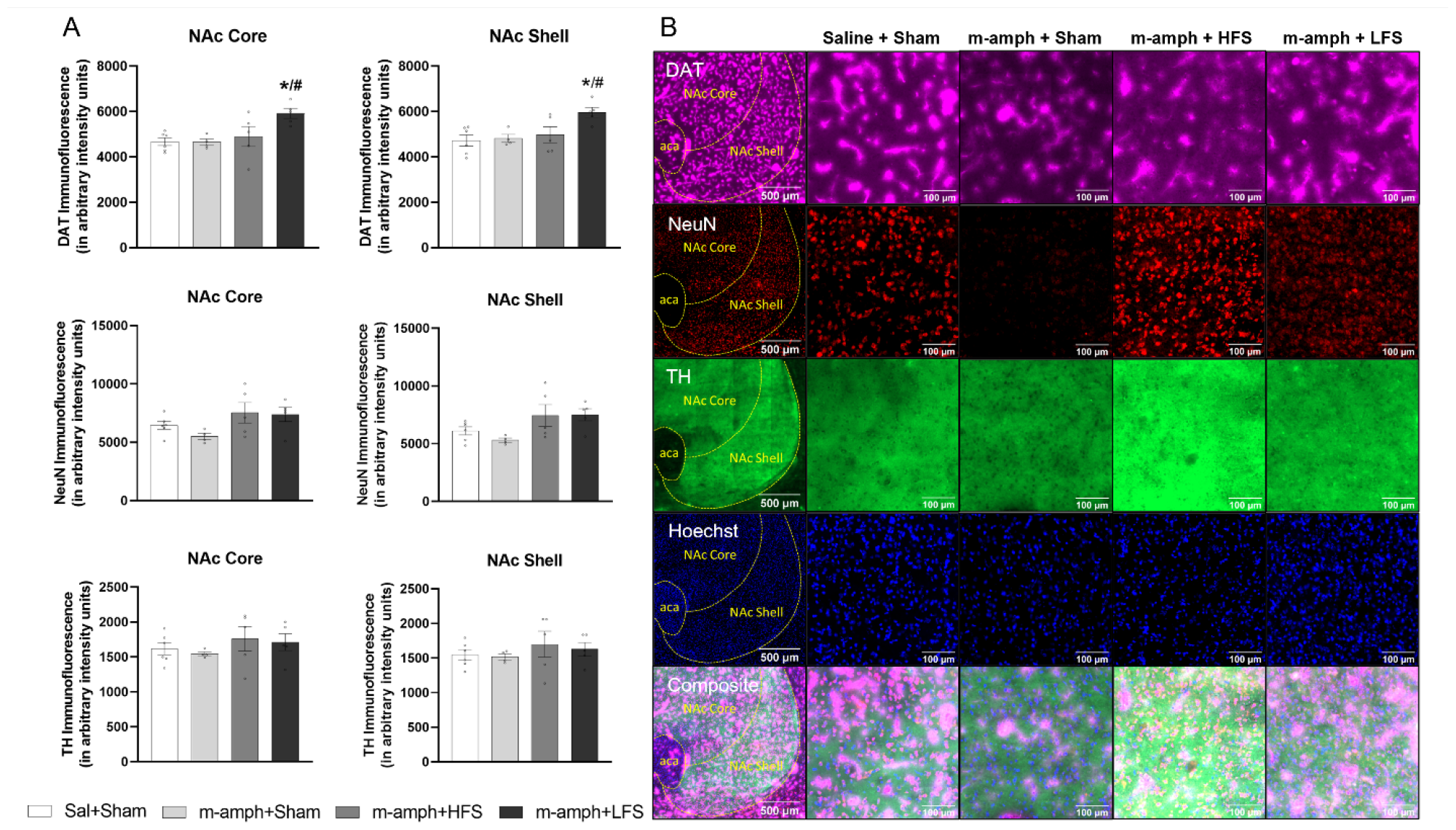
Effects of high frequency stimulation (HFS) and low frequency stimulation (LFS) on nucleus accumbens (NAc) DAT, NeuN and TH expression in rats submitted to the animal model of mania induced by methamphetamine (m-amph). **(A)** Protein immunofluorescence presented in arbitrary intensity units, bars represent mean and standard error according to one-way ANOVA. *=p<0.05 compared to the Saline+Sham; #=p<0.05 compared to the m-amph+Sham group according to Tukey’s post-hoc test. **(B)** First column represents a sample image NAc region staining, second to fifth columns shown representative images detailing DAT (far red), NeuN (red) and TH (green) staining for each experimental group.

## 4. Discussion

Chronic m-amph injection increased locomotor activity in the OFT when compared to the control group. Hyperlocomotion is considered one of the most objective manic-like behaviors assessed in animals, akin to states of increased activity, energy, or drive in humans (31). Administration of psychostimulants, such as amphetamines, is known for inducing manic-like behaviors, and is considered a reliable animal model of mania, being based on monoaminergic dysregulation which responds to mood-stabilizer treatments, supporting face, construct and predictive validities (23). M-amph treatment increases time spent in the center of the OFT arena, which is considered a parameter of risk-taking behavior and decreased anxiety in rodents, which may be translated to poor judgment associated with mania (32, 33). In the present study, m-amph treatment appeared to increase risk-taking behavior, however this increase was not statistically significant.

Electrode implantation alone was able to reduce hyperactivity induced by chronic m-amph, although it did not alter locomotion in saline-treated animals. The positive effects of sham DBS are described in several depression clinical trials, although the effects of DBS are more substantial (34, 35). Perez-Caballero and colleagues (36) described that sham-DBS antidepressant effect is related to increased expression of inflammatory mediators and activation of several short-term repair mechanisms in the region of electrode implantation which can last for up to 2 weeks after electrode implantation and is blocked by anti-inflammatory drugs. Although not described in mania models so far, activation of inflammatory mediators and repair mechanisms in the VTA might contribute to anti-manic-like effects observed in the sham group. Since electrode implantation did not change behavior in saline animals it is unlikely that reduced locomotion observed in Sham/DBS groups is due VTA lesioning-induced locomotion impairment in m-amph group.

Both LFS and HFS were able to completely reverse m-amph-induced hyperlocomotion. Studies describing the effects of DBS on m-amph-induced hyperlocomotion are scant. However, Perez et al. (37) demonstrated that hippocampal HFS DBS reversed hyperactivity induced by acute m-amph administration in sensitized animals in a model of schizophrenia. DBS-induced inhibition of the hippocampus indirectly decreases VTA activity and dopaminergic signalling in the NAc, normalizing the behavioral responses (37). Those results are consistent with the present study, suggesting that direct inhibition of VTA through LFS and HFS DBS normalize m-amph induced hyperlocomotion.

We further investigated the role of phasic DA release on the behavioral effects of VTA LFS and HFS DBS in a different set of animals. For the saline-treated group, phasic DA release and reuptake time did not change post-sham when compared to baseline. Although not related to the first experiment protocol, experiment 2 animals received a m-amph challenge after the post-sham period to confirm DA responses. As previously described by other authors, an acute m-amph injection increased both evoked DA release and reuptake time (38). M-amph increases DA release through different mechanisms such as disruption of the dopamine transporter (DAT) (39), and increased DA synthesis after sub-toxic doses (<2.5mg/kg) (40, 41).

Like the saline group, m-amph-treated animals did not show any significant difference between evoked DA post-sham to baseline. Interestingly, time to DA reuptake was increased after sham when compared to baseline in this group. To the best of our knowledge this is the first study showing late effects of a m-amph challenge on phasic DA reuptake in pre-sensitized animals, considering baseline recordings were acquired 100min after m-amph injection and post sham acquired 120min after injection. The mechanisms underlying this effect remain unknown, however a previous study demonstrated that high extracellular concentrations of DA after phasic bursts can mediate internalization of DATs. As a result, upon cessation of phasic bursts, extrasynaptic DA clearance is slower, thus accumulating in the extracellular space (43). This rebound mechanism could explain, at least in part, the late effect of mamph on phasic DA dynamics observed after the sham stimulation period.

HFS and LFS had similar effects on phasic DA dynamics. Although only LFS was able to decrease phasic DA release, both patterns of stimulation prevented the slower DA reuptake observed in the m-amph+sham group. The effects of intermittent LFS on DA release remain unknown, however this programmed LFS was designed to mimic the natural VTA firing configuration of a healthy rat, being able to normalize BDNF levels and alleviate depressive-like behavior in an animal model of depression (16, 44). Although it’s described that the hyperdopaminergic activity observed in *ClockΔ19* mutant mice, a genetic model of mania, is due to the increased firing rate of dopaminergic neurons in the VTA (22), little is known about the pattern of VTA firing in chronic m-amph treated animals. *In vitro* studies have demonstrated that low doses of methamphetamine can enhance dopamine neurotransmission and increase dopamine neuron firing through a DAT-mediated excitatory conductance (45). Thus, it is possible that chronic m-amph treatment alters VTA dopaminergic neurons firing in animals in a DAT-dependent manner, which is corrected by LFS.

In fact, LFS increased NAc DAT expression, suggesting that its effects on phasic DA may be driven by DAT upregulation. In this regard, Miller and colleagues (46) demonstrated that VTA dopaminergic network connectivity is regulatable via the DAT but not a D2 receptor dependent mechanism, suggesting a master regulator role of this transporter. There is a genetic report suggesting linkage disequilibrium between bipolar disorder and the dopamine transporter (47). A PET neuroimaging study showed lower caudate transporter availability in bipolar disorder than in controls (48). Zhao et al. (49) demonstrated that chronic HFS can increase DAT levels, playing an important role on dopaminergic system modulation in an animal model of Tourette syndrome. Interestingly, a case report also demonstrated that 1 year of NAc HFS was able to increase striatal levels of DAT in an m-amph use disorder patient (50). Together with literature data, the present study suggests that DAT upregulation may act to re-establish normal DA dynamics, playing a key role on the anti-manic effects of DBS.

It is important to note that some limitations of the present study should be addressed in future investigations. The OFT test was conducted immediately after the VTA DBS session. Simultaneous DBS stimulation and OFT performance may provide a better temporal resolution of the behavioral effects of LFS and HFS VTA DBS. Animals submitted to the experiment 2 design were under urethane anaesthesia during phasic DA release recordings, which could affect the comparison of behavioral and electrochemical analysis. FSCV analysis in awake animals (even during OFT) might better reflect DA dynamics and mania-like behavior. FSCV is an electrochemical technique developed to measure rapid changes in neurotransmitters levels, and since it uses the method of background subtraction it can only measure phasic DA release but not tonic levels (51). Evaluation of tonic DA levels (through micro dialysis or multiple-cyclic square wave voltammetry (52)), and DAT activity would help to better understand the role of DA dynamics underlying the potential anti-manic effects of LFS and HFS VTA DBS.

## 5. Conclusion

In summary, this study demonstrated that both LFS and HFS acute DBS of the VTA region exerts anti-manic effects in animals with m-amph induced mania. In addition, LFS and HFS modulated DA dynamics (more specifically DA reuptake) and DAT expression in the NAc, suggest a potential mechanism whereby VTA DBS may exert its anti-manic effects. The next step is to better understand the real-time effects of DBS in manic-like behavior, as well as other aspects of DA signalling. However, these results point to the potential application of neuromodulation approaches to model and clinically modulate manic-like behaviours.

## Acknowledgments

Funding for this project was provided by the National Health and Medical Research Council (NHMRC; APP1160472) (Australia) *Coordenação de Aperfeiçoamento de Pessoal de Nível Superior* (CAPES) (Brazil). MB is supported by a NHMRC Senior Principal Research Fellowship and Leadership 3 Investigator grant (GNT1156072 and 2017131). RV is supported by CREDIT: The CRE for the Development of Innovative Therapies for Psychiatric Disorders (GNT1153607). The authors gratefully acknowledge the QBI Advanced Microscopy and Histology facilities for their support and assistance in this work.

