## Supplementary figure 1 for "Anti-manic effect of deep brain stimulation of the ventral tegmental area in an animal model of mania induced by methamphetamine"

### Supplementary material

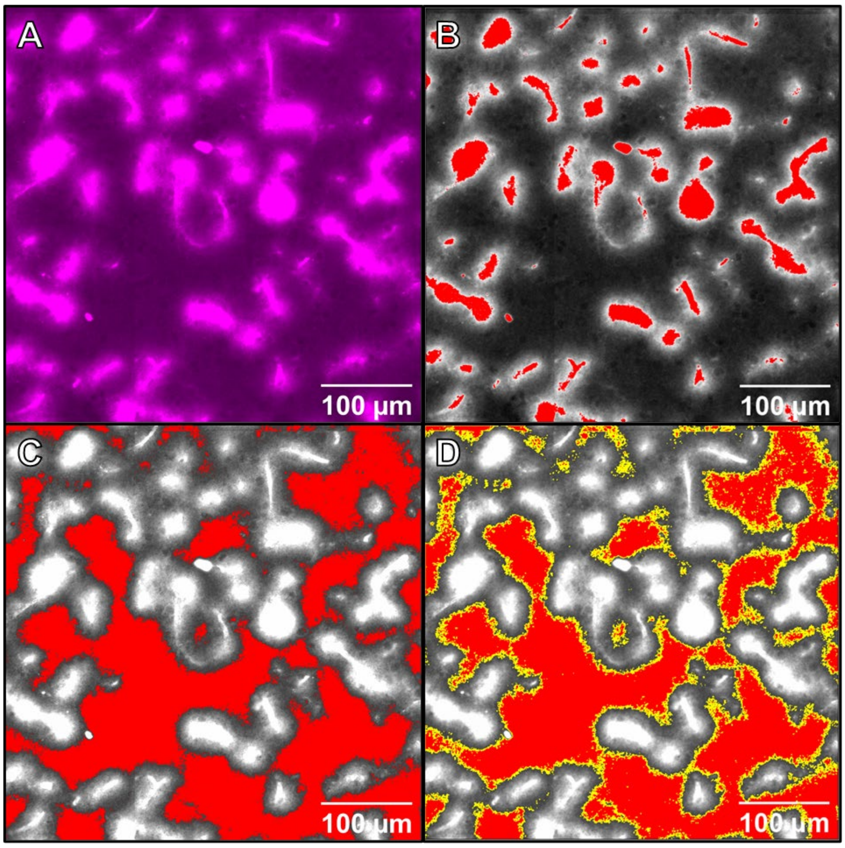


**Supplementary Figure 1:** Step-by-step representation of out effect correction prior to semi-quantitative intensity analysis. (A) Raw dopamine transporter (DAT) staining sample image; (B) Detection of blood vessel intensity; (C) Intensity selected threshold range to exclude DAT positive blood vessels and surrounding area; (D) Selected area for intensity analysis.
